# First detection of Infectious Spleen and kidney Necrosis Virus (ISKNV) associated with massive mortalities in farmed tilapia in Africa

**DOI:** 10.1101/680538

**Authors:** José Gustavo Ramírez-Paredez, Richard K. Paley, William Hunt, Stephen W. Feist, David M. Stone, Terence R. Field, David J. Haydon, Peter A. Ziddah, Mary Nkansa, Emanuel K. Pecku, Joseph A. Awuni, James Guilder, Joshua Gray, Samuel Duodu, Timothy S. Wallis, David W. Verner-Jeffreys

## Abstract

In late 2018, unusual patterns of very high mortality (>50% production) were reported in intensive tilapia cage culture systems across Lake Volta in Ghana. Samples of fish and fry were collected and analysed from two affected farms between October 2018 and February 2019. Affected fish showed darkening, erratic swimming and abdominal distension with associated ascites. Histopathological observations of tissues taken from moribund fish at different farms revealed lesions indicative of viral infection. These included haematopoietic cell nuclear and cytoplasmic pleomorphism with marginalisation of chromatin and fine granulation. Transmission electron microscopy showed cells contained conspicuous virions with typical Iridovirus morphology i.e. enveloped, with icosahedral and or polyhedral geometries and with a diameter c.160 nm. PCR confirmation and DNA sequencing identified the virions as Infectious Spleen and Kidney Necrosis Virus (ISKNV). Samples of fry and older animals were all strongly positive for the presence of the virus by qPCR. All samples tested negative for TiLV and Nodavirus by qPCR. All samples collected from farms prior to the mortality event were negative for ISKNV. Follow up testing of fish and fry sampled from 5 additional sites in July 2019 showed all farms had fish that were PCR positive for ISKNV, whether there was active disease on the farm or not, demonstrating the disease was endemic to farms all over Lake Volta by that point. The results suggest that ISKNV was the cause of disease on the investigated farms and likely had a primary role in the mortality events. A common observation of coinfections with *Streptococcus agalactiae* and other tilapia bacterial pathogens further suggests that these may interact to cause severe pathology, particularly in larger fish. Results demonstrate that there are a range of potential threats to the sustainability of tilapia aquaculture that need to be guarded against.

## Introduction

The farming of tilapia species (*Oreochromis* spp.) is one of the most important sectors in aquaculture worldwide with total global production estimated at more than 6 686 000 tonnes in 2016 (FAO, 2018). In Africa, production is still dominated by Egypt with more than 1 000 000 tonnes produced in 2017 (1). However, tilapia culture has become increasingly important in several other African countries, where it boosts the local economy and constitutes an affordable source of animal protein for human consumption.

In Ghana, Nile tilapia production in 2016 had reached more than 50 000 tonnes, from only 2 000 tonnes per year in 2006 (FAO, 2016) with more than 90% of the production derived from high stocking density floating cage systems in Lake Volta. However, as production systems have intensified in the area, the industry has been increasingly affected by a range of disease issues (Jansen, Cudjoe, & Brun, 2018; Verner-Jeffreys et al., 2018).

In 2017 (Verner-Jeffreys et al., 2018) conducted the first comprehensive disease investigation in tilapia farmed in Lake Volta Ghana. *Streptococcus agalactiae* multilocus sequence type 261 was shown to be a major cause of mortality for farmed Nile tilapia and a range of other bacterial and parasitic pathogens including *Aeromonas* spp., *Streptococcus iniae, Flavobacterium columnare* and a *Myxobolus* sp. were also detected.

Jansen et al. (Jansen et al., 2018) conducted follow up studies, including a broad-ranging epidemiological investigation and suggested that mortalities caused by bacteria in Lake Volta were not a major concern for the local economy as farmers had managed to sustain losses by increasing production of fingerlings, treatment with antibiotics and use of autogenous vaccines.

Historically, bacterial infections were the major threat for the health of farmed tilapia (Plumb & Hanson, 2011). However, in recent years a number of viral diseases have emerged worldwide with devastating effects for the industry(Machimbirike et al., 2019). Tilapia Lake Virus (TiLV) is considered the main viral challenge to tackle, as it has spread to many producing countries, causing high mortalities in all production stages. Although TiLV has been detected elsewhere in Africa (Hounmanou et al., 2018), the initial study by (Verner-Jeffreys et al., 2018), diagnostic investigations undertaken by Ridgeway Biologicals Ltd. (Ramirez *et al*. unpublished data), or the more recent survey undertaken by (Jansen et al., 2018), all failed to provide any evidence of TiLV in diseased tilapia reared in Lake Volta. Up until September 2018, with the exception of the detection of a nodavirus sequence by (Verner-Jeffreys et al., 2018), no significant association between viral agents and tilapia mortality events have been demonstrated in Ghana.

From September 2018 to March 2019 outbreaks of disease with very high levels of morbidity and mortality (60-90%), were experienced in both vaccinated and unvaccinated tilapia by farmers in Lake Volta. In late-September 2018, a farm located below the lower dam in the region of Asutsuare, was the first to suffer episodes of massive acute mortalities (https://goo.gl/LmqbG2 and https://bit.ly/2NwDEbD). Approximately a week after the first report, a second farm located in the Akuse region (~5 km upstream of Asutsuare, but still below the lower dam) also experienced acute mortalities. By mid-October, multiple floating cage-based farms in the Dodi region (above the upper dam) reported losses of more than 10 tonnes per day https://goo.gl/yj4oT4. In late November, farmers that had been unaffected in the Asikuma region (downstream of Dodi but still above the upper dam) also started to suffer episodes of massive acute mortality. (**Figure 1**).

**Figure 1.**
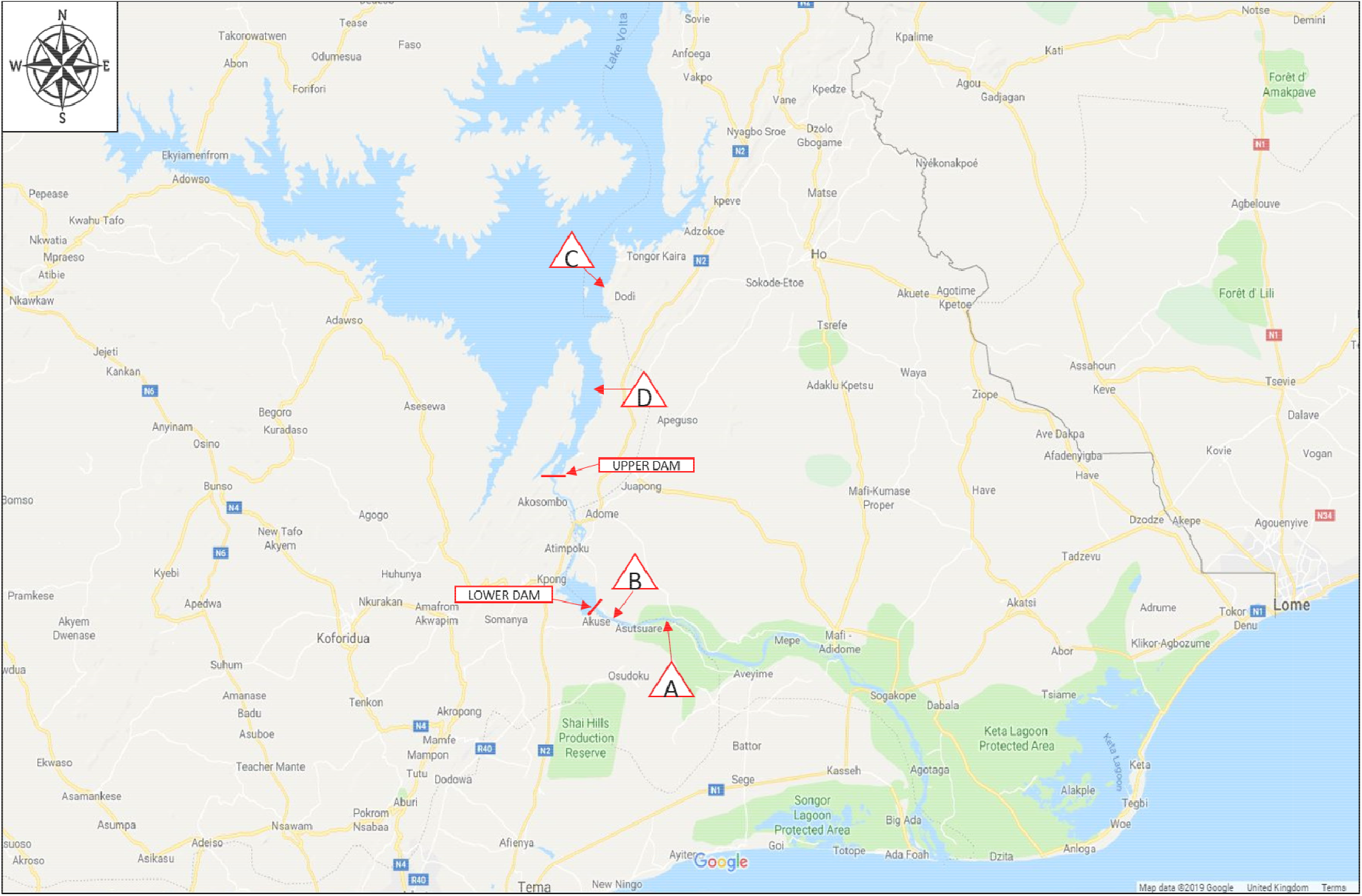
Map of lower region of Lake Volta in Ghana, West Africa. Red triangles indicate the regions in chronological order (A to D) where the outbreaks of mortality occurred.

By the end of 2018, most tilapia farmers in Lake Volta had reported mortalities that they were not able to contain by increased production of fingerlings or treating with antibiotics (**Supplementary File 1**, https://bit.ly/2KZFFuw and https://goo.gl/cPmpSE). Mortality events continued into and throughout 2019. We report the results of a comprehensive disease investigation, conducted at seven affected farms from two different regions of Lake Volta, to gain insights into the causes of these mortalities.

## 2 Materials and Methods

### 2.1 Sampling

Farm 1, a medium size (approximately 600-800 tonnes per annum production) cage culture operation on upper Lake Volta, was visited on the 18/10/18. Samples from 10 fish (average weight 200g) were taken for bacteriology. In addition five sets of samples containing liver and brain from fish from different cages were collected for molecular diagnostics as detailed in **Table 1**.

**Table 1.** Summary of sampling and results

| Sampling date & Farm | No. of Fish and size | Farm observations and ‡clinical signs | Bacteriology | Histopathology | Virology Results<sup>¥</sup><br>[ ] – qPCR<sub>RSIV/ISKNV</sub> copy number/sample reaction range |
| --- | --- | --- | --- | --- | --- |
| 18/10/2018<br>†Farm 1 | n=5; 100-300g | Wide scale ongoing mortalities in on growing fish in cages. | *Aeromonas veronii*;<br>*Aeromonas jandaei*;<br>*Plesiomonas shigelloides*;<br>*Chryseobacterium sp*;<br>*Acinetobacter johnsonii*; | Not done | *qPCR<sub>TILV</sub> 5/5 -ve<br>*cPCR<sub>a</sub> ISKNV 4/5 +ve<br>*qPCR<sub>RSIV/ISKNV</sub> 5/5 +ve<br>[7.5 x 10<sup>1</sup> – 4.60 x 10<sup>5</sup>] |
| 28/11/2018<br>†Farm 2 | n=14; 74-401g | Wide scale ongoing mortalities in on growing fish in cages (**Figure 2**). On shore fry production unit: no unusual mortalities. | *Aeromonas veronii*;<br>*Aeromonas jandaei*;<br>*Streptococcus agalactiae* capsular type Ib biotype 2 (non-haemolytic);<br>*Edwardsiella tarda* | Evidence of Gram negative and Gram positive bacterial and viral# infection (including in same fish) | qPCR<sub>TILV</sub> 14/14 -ve<br>qPCR<sub>NODA</sub> 14/14 -ve<br>*cPCR<sub>a</sub> ISKNV 12/14 +ve<br>*qPCR<sub>RSIV/ISKNV</sub> 14/14 +ve<br>[2.48 x 10<sup>1</sup> - 3.3 x 10<sup>6</sup>] |
| 17/12/2018<br>†Farm 2 | n=7; 40-646g | Much lower mortalities in on growing fish in cages than previous sampling visit. On shore fry production unit: no unusual mortalities. | Not done | Evidence of bacterial infection but not of viral infection | qPCR<sub>TILV</sub> 7/7 -ve<br>qPCR<sub>NODA</sub> 7/7 -ve<br>**cPCR<sub>b</sub> RSIV/ISKNV 1/7 +ve**<br>**megalocytivirus, sequence = ISKNV**<br>*qPCR<sub>RSIV/ISKNV</sub> 6/7 +ve<br>[1 x 10<sup>0</sup> - 24 x10<sup>0</sup>] |
| 20/02/2019<br>†Farm 2 | n=14; 6-9 cms | Limited mortalities in on growing fish in cages. Very severe mortalities in on shore fry production unit. | Not done | Very severe virus-associated pathology (in tissue samples from all fish). | qPCR<sub>TILV</sub> not done<br>qPCR<sub>NODA</sub> not done<br>cPCR<sub>a</sub> ISKNV 7/7 +ve<br>qPCR<sub>RSIV/ISKNV</sub> 13/13 +ve<br>[5.1 x 10<sup>5</sup> – 1.51 x 10<sup>7</sup>] |
| 09/07/2019<br>Farm 3 | Grp 1, n=3; 60-80g<br>Grp 2, n=3; 20-40g<br>Grp 3, n=5; 1-2g fry<br>Grp 4, n=5 pools of ≤0.2g fry | No mortality events reported at site. Grp 1 on-growing in lake, Grps 2-4 in ponds. | Grps 1 & 2 - No significant colonies<br>Grps 3-5 Not done | Evidence of metacercaria, bacterial gill epitheliocystis and myxosporidiosis in Grps 1 & 2. - No evidence of viral infection. | cPCR<sub>b</sub> RSIV/ISKNV<br>Grp 1 - 2/3 +ve (nested only)<br>Grp 2 - 3/3 +ve (nested only)<br>Grp 3 – 5/10 +ve (nested only)<br>Grp 4 - 3/10 +ve (nested only) |
| 09/07/2019<br>Farm 4 | n=5 pools of<br>≤0.2g fry | Reported recent mortality<br>event now recovering. No<br>clinical signs | Not done | Myxosporidian cysts<br>in cranial sub-<br>epithelial and<br>connective tissue | cPCRb<sub>RSIV/ISKNV</sub> 2/5 +ve (nested only and<br>in one of duplicates only) |
| 10/07/2019<br>Farm 5 | n=5 x3 fish organ<br>pools; ~40g | Ongrowing fish on lake,<br>recent mortality event in<br>adjacent cage cleared<br>previous month, no clinical<br>signs in sampled fish | No significant colonies | not done | cPCRb<sub>RSIV/ISKNV</sub> 5/5 +ve (nested only,<br>one of which weak in 1st round) |
| 10/07/2019<br>Farm 6 | n=10 x3 fish<br>pools; 0.2-0.75g | Ongoing losses of approx.<br>30% in fry pools. Fry<br>observed with distended<br>abdomen and disturbed<br>swimming. | Not done | Evidence of viral<br>infection,<br>occasionally severe | cPCRb<sub>RSIV/ISKNV</sub> 10/10 +ve<br>Virus isolation – BF cells |
| 10/07/2019<br>Farm 7 | n=5 x2 fish organ<br>pools; 20-40g | Internal haemorrhaging,<br>ascites, pale kidney, anaemia,<br>reduced spleen and enlarged<br>liver | No significant colonies | Evidence of viral<br>infection | cPCRb<sub>RSIV/ISKNV</sub> 5/5 +ve<br>Virus isolation – BF cells |

† See Materials and Methods for farm descriptions
‡ All moribund fish, including fry, presented similar signs: erratic swimming, lethargy, ascites, swollen, dark spleen, haemorrhagic livers and other organs.

Farm 2, a large cage culture operation (>2000 tonnes per annum), was visited for sampling on 28/11/18, 17/12/18 and 20/02/19. For the first two visits, moribund fish between 40-646g collected from cages on the main lake were sampled. In the first visit for bacteriology and virology on the second visit for virology only. For the third visit, responding to reports that they were now experiencing very heavy mortalities in their fry production units (>70%), samples of moribund fry and juveniles from both nursery cages on the main lake and from their onshore hatchery supplied with water pumped from the main lake, were analysed for virology. From this farm material was also taken for histological and molecular diagnostic investigations during the visits as detailed in **Table 1 and Supplementary File 2**. During the visits, semi-structured interviews were carried out with farm managers and/or workers. Interview questions were primarily constructed to ascertain trends in mortality levels since September 2019, any observed clinical signs in this time and whether there were any differences in impact associated with fish life stage and/or system. Additionally, relevant information on potential risk factors, mitigation measures and biosecurity practices, as well as any further farmer concerns, were discussed.

A further 5 farms in the Akosombo, Atimpoku and Dasasi regions were visited and sampled for virology, bacteriology, histology and molecular diagnostic investigation from 9-10/07/2019. These included fry, on-growing and broodstock fish from farms of varying capacity and with either no reported disease, fish that had survived recent mortality events on farms or fish with ongoing clinical signs or mortalities.

All moribund fish from the visited farms were humanely euthanised with a lethal overdose of tricaine methanesulfonate 1,000 mg/g (Pharmaq, Hampshire, UK) followed by brain destruction prior to the necropsy.

### 2.2 Bacteriology

Samples for bacteriology were collected from the brain, liver, kidney and spleen with sterile cotton swabs and inoculated onto tryptone soya agar (TSA), Columbia blood agar (CBA), Tryptone yeast extract salts agar (TYES) (Southern Group Laboratory, Corby, UK) and cystine heart agar with 2% bovine haemoglobin (CHAH) (Becton Dickinson, Oxford, UK).

All inoculated agar plates were incubated at 28 °C for 24-72 hours. Colonies assessed as significant based on occurrence and dominance were subcultured to purity on similar media. Pure relevant isolates were initially identified by morphology and Gram staining. The partial 16S rRNA genes of the Gram negative isolates identified were PCR amplified and sequenced using the method described by (Klindworth et al., 2013). Gram positive cocci forming chains were screened using a *Streptococcus agalactiae* specific capsular typing multiplex PCR developed by (Shoemaker, Xu, Garcia, & Lafrentz, 2017). The Gram negative isolates strains confirmed as *Aeromonas* spp. based on partial 16S rRNA gene sequence analysis were further characterised based on partial *gyrB* sequencing analysis for identification at the species level as described by (Yáñez, Catalán, Apráiz, Figueras, & Martínez-Murcia, 2013).

### 2.3 Histopathology and Electron Microscopy

Tissues were fixed in neutral buffered formalin (NBF) for a minimum of 24 hr before being placed in glycerol diluted 50:50 with phosphate buffered saline (PBS) for transportation to Cefas. On receipt, tissues were rinsed in 70% alcohol and placed again in NBF for a final period of fixation prior to processing using standard protocols in a vacuum infiltration processor. Tissue sections were cut at a thickness of 3-4 μm on a Finnese^®^ microtome, mounted on glass slides and stained with haematoxylin and eosin using an automated staining protocol. Stained sections were examined for general histopathology by light microscopy (Nikon Eclipse E800). Digital images and measurements were obtained using the Lucia^™^ Screen Measurement software system (Nikon, UK).

For electron microscopy, small samples of tissues fixed in NBF as above were rinsed three changes of 0.1 M sodium cacodylate buffer, followed by post fixation in 2.5% glutaraldehyde in the same buffer for 1 hour prior to a second post fixation for 1 hour in 1 % osmium tetroxide in 0.1 M sodium cacodylate buffer. Subsequently, fixed tissues were dehydrated in an ascending acetone series acetone series and embedded in epoxy resin 812 (Agar Scientific pre-Mix Kit 812, Agar scientific, UK) and polymerised at 60 °C overnight. Semi-thin (1 μm) survey sections were stained with 1 % Toluidine Blue and examined by light microscope to identify areas of interest. Ultrathin sections (70-90 nm) of the targeted areas were placed on uncoated copper grids and stained with uranyl acetate and Reynold’s lead citrate (Reynolds 1963). Grids were examined using a JEOL JEM 1400 transmission electron microscope and digital images captured using a GATAN Erlangshen ES500W camera and Gatan Digital Micrograph^™^ software.

### 2.4 Molecular diagnosis (viral and bacterial detection)

The samples collected for molecular diagnosis were washed twice in 750μl of sterile 1x PBS to remove the RNA-*later*^®^ and homogenised. Total nucleic acids were extracted using nanomagnetic beads i.e. Genesig Easy DNA/RNA Extraction Kit (Primerdesign, Southampton, UK) and stored until further use.

#### *Multiplex PCR for detection of Streptococcus* spp

Nucleic acids extracted were used as a template on a multiplex PCR (unpublished data) to confirm the presence of *Streptococcus* spp. that had been previously reported as fish pathogens including *Streptococcus agalactiae* (Delannoy et al., 2013), *Streptococcus uberis* (Luo et al., 2017) and *Streptococcus dysgalactiae* (Abdelsalam, Asheg, & Eissa, 2013).

#### qPCR for detection of TiLV and Nodavirus

Nucleic acids were used for the detection of tilapia lake virus and nodavirus by quantitative PCR using the commercial kits: Path-TiLV-EASY and Path-Betanodavirus-EASY (Primerdesign, Southampton, UK) in the platform Genesig q16^®^ (Primerdesign, Southampton, UK) as per the protocols suggested by the manufacturer.

#### Conventional PCR for detection of megalocytiviruses

The generic PCR protocol for notifiable aquatic megalocytiviruses (Red Seabream Iridoviral disease / Infection spleen and kidney necrosis virus (J Kurita, Nakajima, Hirono, & Aoki, 1998; World Organisation for Animal Health OIE, 2018)) was initially used, to screen the samples of fish collected from visit 2 at Farm 2. For this, genomic DNA was extracted as follows: the RNA-*later*^®^ was removed and the tissue samples weighed. Depending on the weight of the tissue available the samples were diluted in RTL buffer (Qiagen) to provide either a 1:10 w/v or a 1:5 w/v and homogenised per fish i.e. all the organs of each fish into an individual pool using Matrix A and the FastPrep apparatus (MP Biomedicals). Following homogenisation, the samples were diluted further with RTL buffer to give a 1:60 w/v homogenate and total nucleic acid was extracted from 300 μl of the clarified sample using the RNA tissue mini kit without DNase (Qiagen) and eluted in a 60 μl volume.

RT was performed at 37°C for 1 h in a 20 μl volume consisting of 1× M-MLV RT reaction buffer (50 mM) Tris pH 8.3, 75 mM KCl, 10 mM DTT, 3 mM MgCl2) containing 1 mM dNTP, 100 pmol random primers, 20 U M-MLV reverse transcriptase (Promega, Southampton UK) and 4μl of the nucleic acid extracted above.

PCR was performed in duplicate in a 50 μl reaction volume with 2.5 μl of cDNA of total nucleic acid, 25 mM dNTPs, 1 x GoTaq^®^ buffer (2.5 mM MgCl2 solution), 5 pmol of each primer (C1105 5’-GGTTCATCGACATCTCCGCG-3’ and C1106 5’-AGGTCGCTGCGCATGCCAATC-3’) and 1.25 units of GoTaq^®^ DNA polymerase (Promega). The cycling conditions were as follows: 40 temperature cycles (1 min at 95°C, 1 min at 55°C and 1 min at 72°C) after an initial denaturing step (5 min at 95°C) followed by a final extension step of 10 min at 72°C.

Amplified products were electrophoresed in a 2% (w/v) agarose/TAE (40 mM Tris-acetate, pH 7.2, 1 mM EDTA) gel containing 1.0 μg ml-1 ethidium bromide at 120 v for 30 mins and viewed under UV light.

PCR products were excised from the gel and the DNA was extracted and purified by ethanol precipitation. Both strands of the PCR product were sequenced using the ABI PRISM Big Dye Terminator v3.1 cycle sequencing kit and the same primers used for the amplification. The forward and reverse sequences were aligned and a consensus sequence generated using the CLC software (Qiagen). Generated consensus sequences were compared with sequences from GenBank using BLASTN (Altschul et al., 1997) and aligned using the MUSCLE application of the MEGA software version 6 (Tamura, Stecher, Peterson, Filipski, & Kumar, 2013).

In addition, the OIE recommended PCR protocol for notifiable aquatic megalocytiviruses (World Organisation for Animal Health OIE, 2018) developed by (J Kurita et al., 1998) was used was used to screen total nucleic acids extracted from the rest of the samples fixed for viral molecular analyses.

In analyses from farms where clinical disease was not observed, a second round PCR using the nested primers C1073 5’-AATGCCGTGACCTACTTTGC-3’ and C1074 5’-GATCTTAACACGCAGCCACA-3’ (15) was employed.

#### qPCR for detection and quantification of megalocytiviruses

The amount of virus present in the samples was also investigated by qPCR. For this the homogenised tissues were subjected to total nucleic acids extraction (~20mg of each organ) using the Genesig Easy DNA/RNA Extraction Kit (Primerdesign) as described earlier. The extracted nucleic acids were tested using the commercial kit Path-ISKNV-EASY (Primerdesign, Southampton, UK) in the platform Genesig q16^®^ (Primerdesign) as per the manufacturer instructions. This detects both red sea bream iridovirus and ISKNV variants. In all cases fish were individually analysed either by pooling liver, spleen and brain or screening individual tissues.

#### Retrospective analyses of archived samples by qPCR

A total of 16 samples of archived tissue homogenates from 5 different farms (that included Farms 1 and 2) were retrospectively screened for ISKNV by qPCR with the commercial kit Path-ISKNV-EASY as described before. From these 7 had been collected during 2017 and the rest in March 2018 (**Supplementary File 5**). All the samples had been previously confirmed as negative for TiLV and Nodavirus using the commercial kits Path-TiLV-EASY and Path-Betanodavirus-EASY.

#### Virus isolation

Frozen spleen and kidney tissue or whole fry of fish showing clinical signs taken from 2 farm sites on Lake Volta on 10 July 2019 were homogenised with sand and pestle and mortar in 1:10 w/v cell culture transport medium (L-15 plus 1% antibiotic antimycotic solution, Gibco). Homogenate was clarified by centrifugation for 10 min at 3000g, inoculated at 1:100 and 1:1000 final dilutions onto GF, BF-2 and E-11 cells in 24 well cell culture plates (Gibco) and incubated at 25°C. After 7 days cells were blind passaged and incubated for a further 7 days. Cells were observed for cytopathic effect (CPE) by light microscopy with phase contrast (IX83 inverted microscope, Olympus, UK).

## 3 Results

### 3.1 Farm visits

On the first visit to Farm 2 on the 27/11/18, farm staff reported very high and ongoing mortalities (Figure 2) in fish bigger than 20g, including broodstock, but no significant losses in fingerlings. Losses reportedly peaked at about 670 crates (equating to approximately 40 tonnes per day) shortly after this visit on the 2/12/2018 **(Figure 2**).

**Figure 2.**
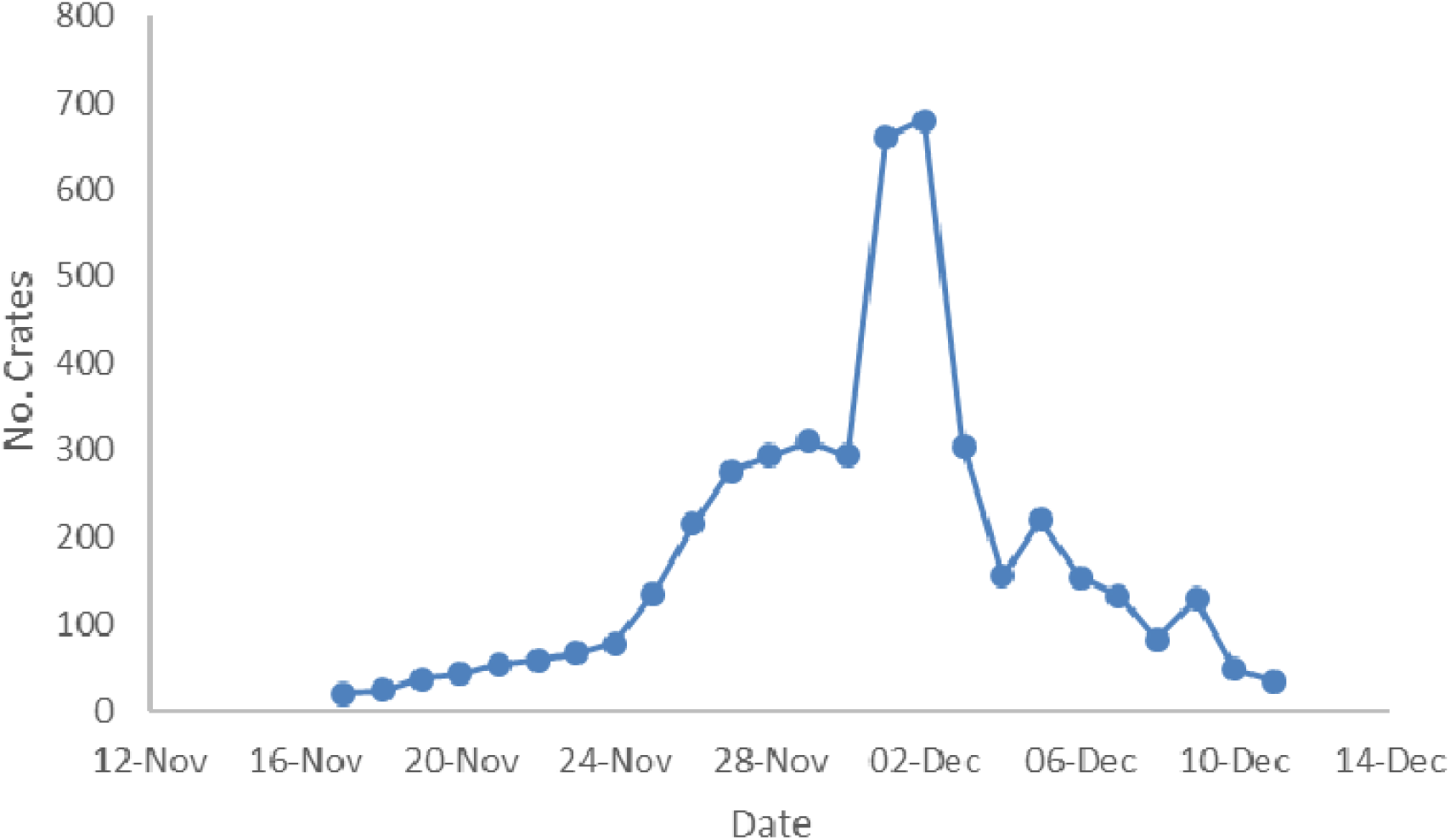
Number of crates of dead/ moribund fish removed from Farm 2 during period of maximum losses. Daily mortalities were estimated based on numbers of crates of rejected ongrowing tilapia collected each day by farm staff. Each crate typically contained approximately 60kg tilapia collected from the cages and rejected because they were either dead, moribund or displayed other adverse signs that prevented their sale.

Losses were so severe that accurate estimation was not possible, with more than 50 additional labourers recruited locally just to remove dead and moribund fish during the peak period. By the second visit to Farm 2, losses of ongrowing fish had reportedly declined back to the background 10-20 crates per day more typically observed i.e. less than 1-2 tonnes per day (**Figure 2**).

During the first two visits to the farms, but particularly to Farm 2, diseased fish were observed swimming away from the school with erratic swimming i.e. on one side, in circles, lethargic, with no equilibrium, upside down etc. **(Supplementary File 3).**

Externally, the fish displayed a range of clinical signs, including skin nodules, frayed fins, loss of eyes, opaque eyes, loss of scales, exophthalmia, anorexia, decolouration or darkened skin, excess of mucous, skin haemorrhages and distended abdomen (**Figure 3)**. At necropsy, fish from both the first visits to Farm 1 and 2 presented with marked ascites, enlarged and haemorrhagic organs including the spleen, heart, brain, gills, but most notably liver and kidney. The gastrointestinal track was empty of solids but contained transparent fluid similar to that also seen in the peritoneal cavity.

**Figure 3.**
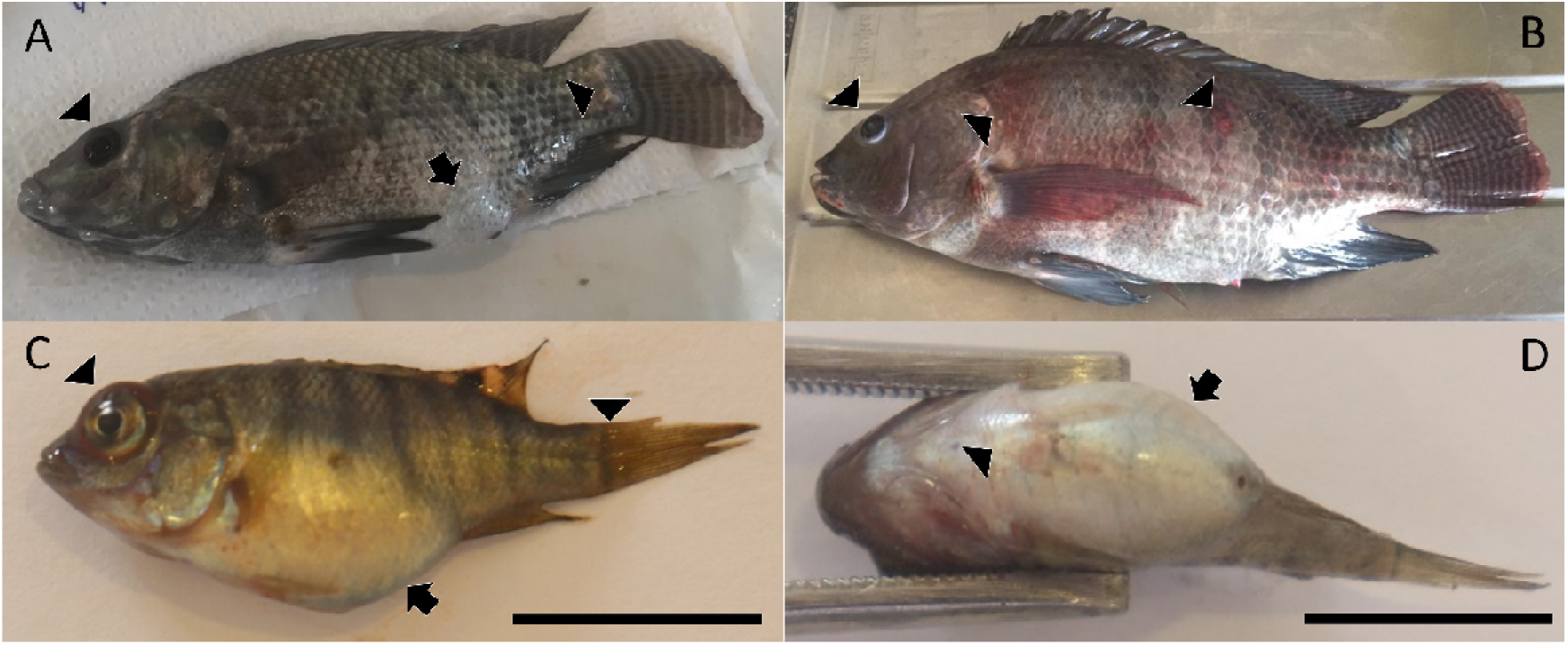
External lesions and clinical signs in diseased Nile tilapia in Lake Volta. **A** Ongrowing fish with emaciation slight ascites (arrow), endophthalmia (left arrow head) and skin purulent abscess (right arrow head). **B** Broodstock with microphthalmia left arrow head), skin haemorrhages (middle arrow head) and skin ulcers (right arrow head). **C** juvenile with exophthalmia (left arrow head), ascites (arrow) and loss of scales, excess of mucus and haemorrhages (right arrow head). Bar = 1cm. **D** Ventral view of juvenile fish presenting severe ascites black arrow and skin haemorrhages (arrow head). Bar= 1cm.

In contrast to the earlier visits, when Farm 2 was visited on 20/02/19, there were very high and ongoing mortalities in the fry production systems (>70%). This was both in their onshore hatchery (supplied with water pumped from the main lake) where eggs were hatched and held until the fry were approximately 20g, and in the nursery cages on the main lake to which fry had been transferred. As with the larger fish sampled, affected fry showed erratic swimming behaviour, skin haemorrhages and severe ascites as the main clinical signs (**Figure 3)**.

### 3.2 *Megalocytivirus* and bacterial infections in Farm 2 during period of high mortalities

Four out of the ten fish examined histologically showed mild tissue necrosis in the spleen and renal haematopoietic tissue with the presence of large numbers of cells showing relative eosinophilia cytoplasmic and nuclear pleomorphism with margination of chromatin in some affected nuclei suggestive of a viral infection diffused throughout the tissue **(Figure 4)**.

**Figure 4.**
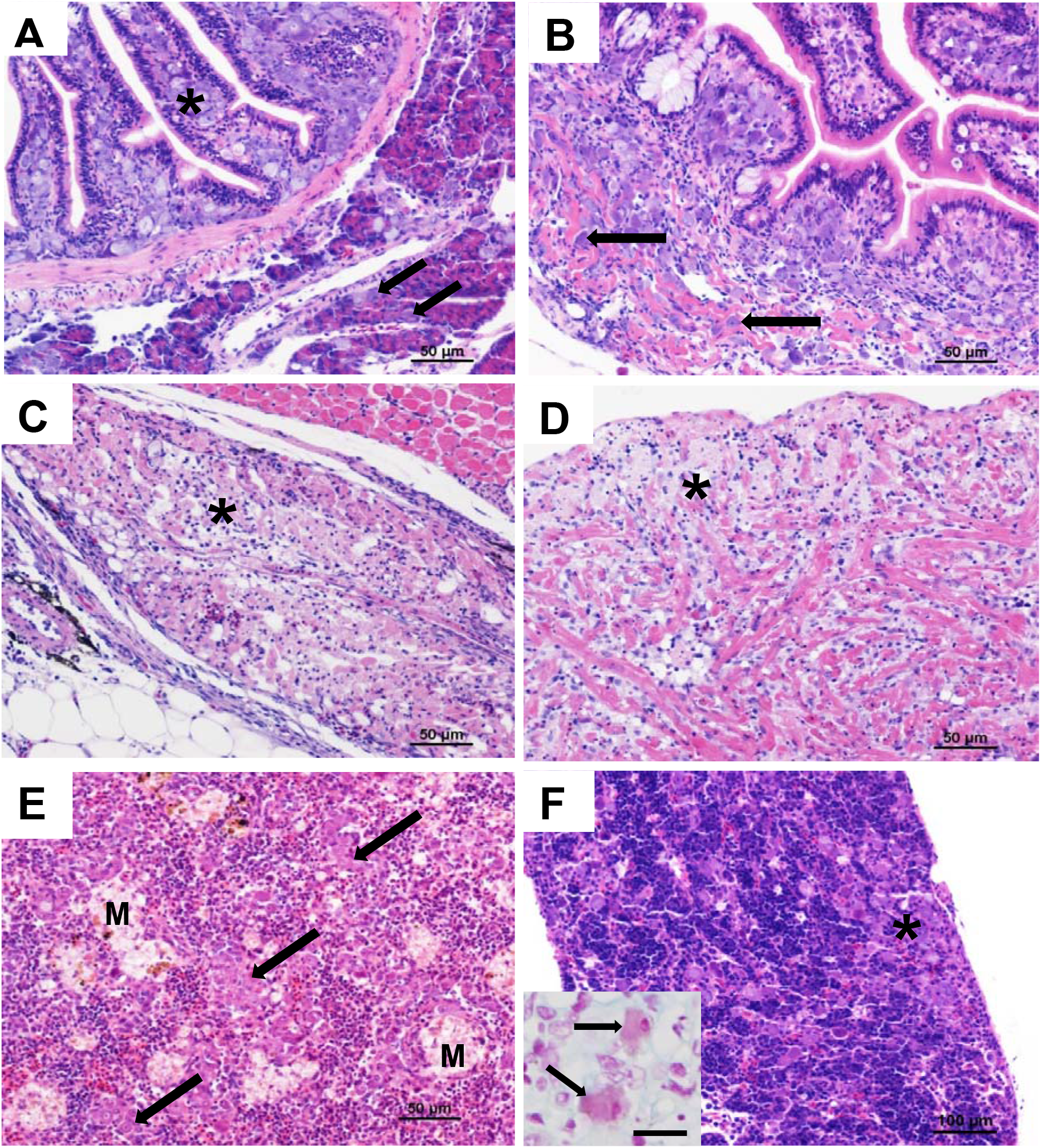
Histopathological cross sections of tissues from diseased Nile tilapia in Lake Volta. All sections stained with H&E unless otherwise stated. **A.** Section through the intestine of an infected fry showing the presence of large numbers of megalocytes in the lamina propria (*) but not affecting the mucosa or underlying muscularis. Note the presence of isolated megalocytes in the pancreatic acinar tissue (arrows). Bar = 50μm. **B.** Same fish as in A, showing infiltration and disruption of the muscularis associated with the presence of megalocytes (arrows). Bar=50 μm. **C.** Pseudobranch showing extensive necrosis (*). Bar=50μm. **D.** Heart showing necrosis of the ventricular muscle (*), particularly in the peripheral regions associated with inflammation. Bar=50μm. **E.** Spleen from a fish from Farm 2 during a mortality episode. Affected cells show pronounced eosinophilia and are distributed throughout the section (*). Numerous vacuolated macrophage aggregates are also present (M). Bar = 50μm. **F.** Pronephros showing extensive distribution of megalocytes without associated tissue necrosis (*). Bar=100μm. Inset shows pale staining of DNA in the cytoplasm of affected cells (arrows). Feulgen stain. Bar=25μm.

The ultrastructure of affected cells revealed the presence of conspicuous viral particles. Some cells showed the presence of numerous diffusely spread virions within the hypertrophied nucleus of affected cells, which also showed degradation of the nuclear membrane (**Figure 5)**. Virions were approximately 160 nm in diameter **(Figure 5D)**. Virion morphology showing icosahedral symmetry with an external double membrane and internal core was consistent with that of viruses from the genus *Megalocytivirus*. In other cases, the nucleus of affected cells appeared condensed and densely stained in histological and resin sections. TEM showed that in these cases the nucleus was tightly packed with virions in various stages of maturation and with some evidence of formation of ‘arrays’ **(Figure 5)**. In two of these cases, a concomitant Gram +ve bacterial infection was also present in the gill and liver. Incidental findings of gill parasites, myxozoan cysts and monogeneans, both present in low numbers as well as low grade epitheliocystis were observed. The brain of a single fish harboured small cysts containing necrotic debris. Gram and Ziehl-Neelsen staining did not demonstrate the presence of bacteria. Other tissues appeared normal. For the set of samples collected at the height of the mortalities from Farm 2 on 28/11/18, *Aeromonas jandaei, Aeromonas veronii* (from skin), *Streptococcus agalactiae* capsular type Ib biotype 2 (non haemolytic) and *Edwardsiella tarda* (from liver and kidney) were recovered.

**Figure 5.**
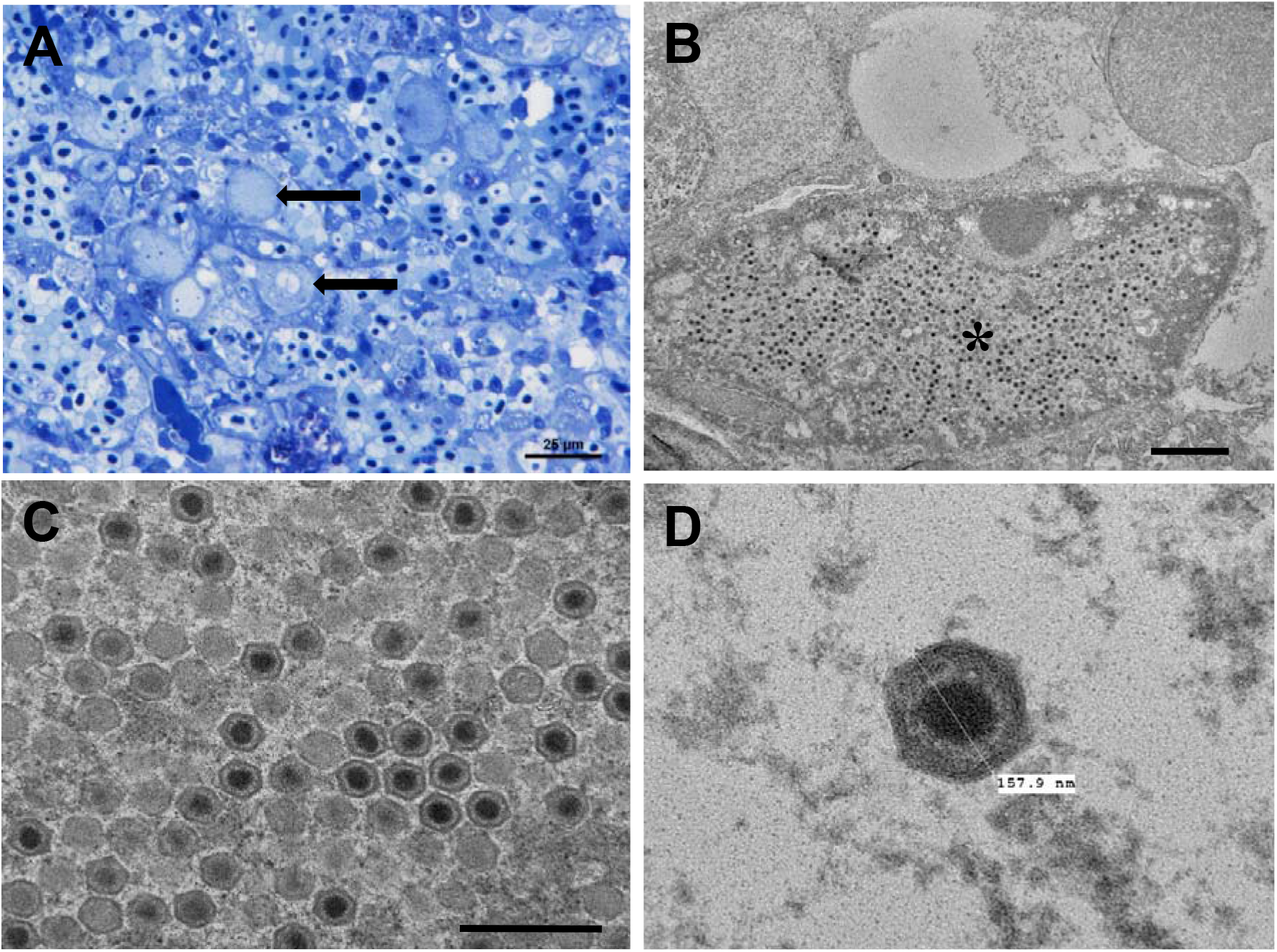
Micrographs of the kidney of diseased Nile tilapia in Lake Volta. **A** Semithin section of affected renal tissue showing characteristic cellular hypertrophy (black arrows). Toluidine Blue stain. Scale bar = 25μm. **B.** Electron micrograph of an individual infected cell with numerous viral particles (*). Adjacent cells appear uninfected. Scale bar **=** 2μm. **C.** Numerous lightly stained developing virions with mature virions. The outer membranes and central electron lucent core are clearly visible. Scale bar = 500nm. **D.** Mature icosahedral virion showing detail of the outer caspid and inner membrane with central electron lucent core.

### 3.3 PCR and sequence confirmation of Megalocytivirus infection

Within the 7 individuals collected from Farm 2 on the second visit that were analysed with the protocol proposed by (Rimmer, A. E., Becker, Tweedie, & Whittington, 2012), a single fish (fish 1) was clearly positive by PCR for RSIV/ISKN and a second very weak product of the correct size was also seen in tissues from Fish 6 (**Supplementary File 4**) The consensus sequence generated from the PCR product from Fish 1 was confirmed as ISKNV sharing 100% nucleotide identity with ISKNV accession no AF371960.1. In the phylogenetic analysis the sequence was assigned to the same lineage as the bulk of the ISKNV sequences (**Figure 6)**.

**Figure 6.**
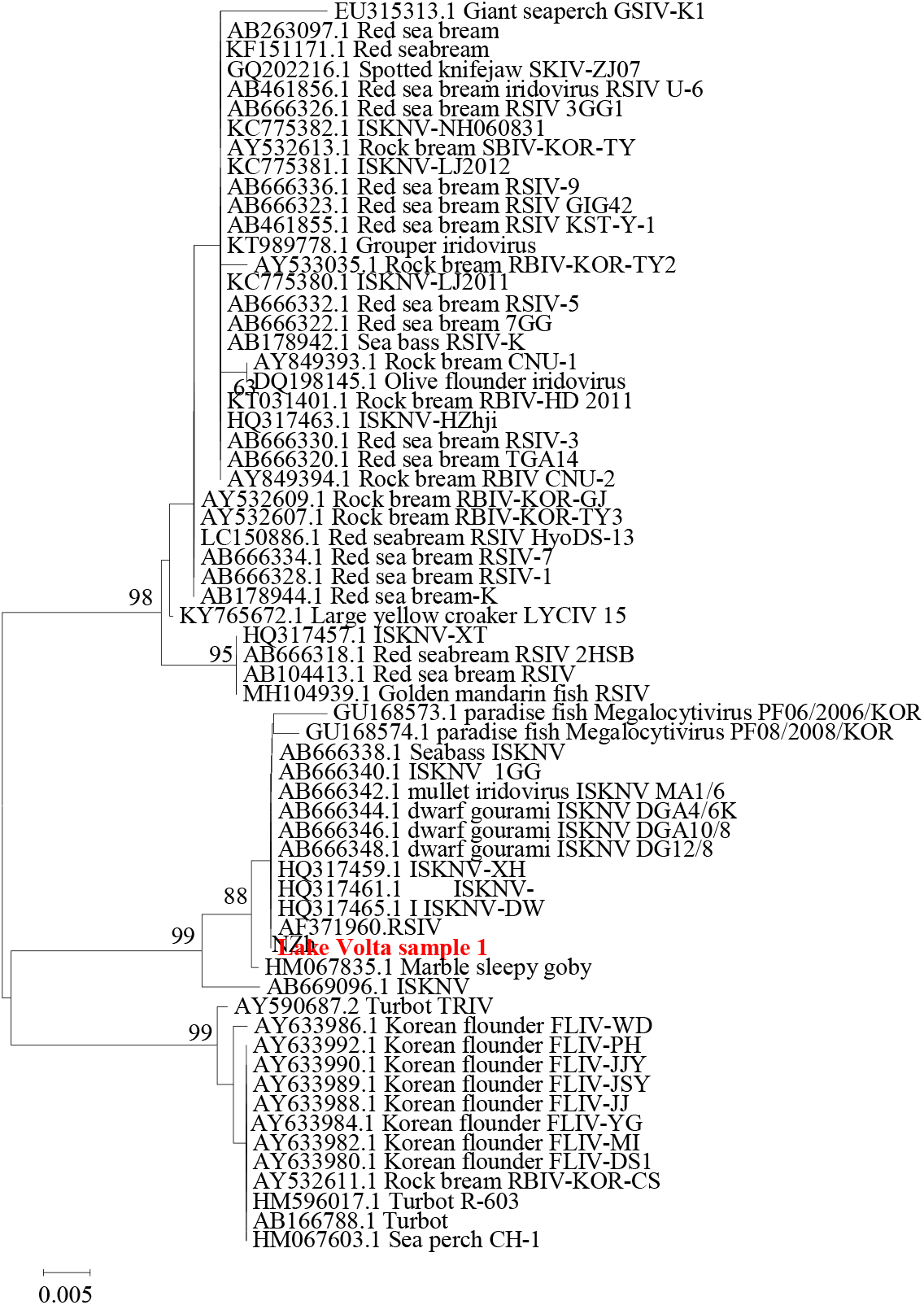
N-J tree showing the genetic relationship of partial MCP sequence from LV#1 to a range of RSIV, ISKNV and related Megalocytiviruses. The GeneBank accession numbers and the host species are included where available.

The samples collected from Farm 1 and Farm 2 at the height of the mortalities were found to be strongly positive when they were retrospectively tested using the (J Kurita et al., 1998; World Organisation for Animal Health OIE, 2018) current OIE recommended PCR method.

### 3.4 qPCR results for megalocytivirus from ongrowing tilapia samples

The qPCR results confirmed that all the fish sampled from Farm 1 in the Akuse region were positive for megalocytivirus. Also, as expected, the samples collected from the first Farm 2 visit during the peak of mortality (28/11/18), were also positive and presented the highest viral titres in grow out fish with some containing over 3 x 10^6^ copies per sample reaction. In contrast, the samples collected from grow out fish during the second visit to Farm 2 had much lower viral copy numbers.

All the archived samples collected in 2017 and March 2018 were negative for ISKNV, when tested by qPCR **(Supplementary File 5)**.

### 3.5 Fry samples had characteristic megalocytivirus pathology and high copy numbers of virus

All the fry samples collected from Farm 2 on 20/02/19, when there were reportedly very high (>90%) losses in that part of their system, were positive for ISKNV by qPCR and these presented the highest titres in the study with some containing up to 1.5 x 10^7^ copies per sample reaction **(Table 1)**. All fry showed moderate to marked histological and pathological features of infection with *megalocytivirus*. Splenic tissues were necrotic and associated with the presence of megalocytes characterised by light sometimes granular cytoplasmic basophilia and hypertrophied nuclei. Kidney also showed the presence of megalocytes but usually with only mild cellular necrosis. The lamina propria in the intestine of a single fish was packed with megalocytes **(Figure 4)**, although necrosis appeared to be absent and the epithelial layer remained intact. Gills showed only minimal focal necrosis, usually affecting the underlying connective tissues. In some fish the choroidal rete was affected with mild necrosis and variable numbers of megalocytes and most fish showed mild myositis with few megalocytes in the skeletal muscle. However, a single fish showed extensive inflammation and myofibrillar necrosis **(Figure 4)**. Connective tissues of the head and in particular around the pharyngeal teeth were often infiltrated with megalocytes. Brain and spinal cord appeared normal. Liver samples were not examined as they were used for virus quantification.

### 3.6 Other pathological observations

*Megalocytivirus*-like pathology was not observed in any of the samples taken from the second visit to Farm 2, two weeks after the peak of mortalities had passed. As with the samples taken at the first visit, there was evidence of bacterial infection in some individuals, particularly fish 5, had marked bacterial infection of the spleen, liver and brain (meningitis). All the samples were positive for the presence of *Streptococcus agalactiae* and negative for *Streptococcus uberis* and S*treptococcus dysgalactiae* by PCR **(Supplementary File 6)**.

A range of different potential bacterial species, including *Aeromonas jandaei* and *Plesiomonas shigelloides* were recovered from fish from Farm 1 **(Table 1)**, but not as pure growths or high quantities, suggesting they had a limited role in observed disease in these animals.

All the samples from Farm 1 and Farm 2 tested for TiLV and from Farm 2 for nodavirus by qPCR were all negative **(Table 1 and Supplementary File 4)**.

The results for all the individual fish tested are shown in Supplementary File 2.

### 3.7 Follow up testing in July 2019

A further 5 farms sampled in July 2019 were tested using nested conventional PCR (15). Fish on farms Farms 6 (fry) and 7 (lake ongrowers) were experiencing ongoing mortality and showing typical clinical signs described above at time of sampling. All samples were strongly positive in a single round assay and virus was isolated in cell culture from both farms. For samples from farms 4 (fry) and 5 (lake ongrowers) which both were recovering from recent mortality events, but had no remaining observable clinical disease, between 40-100% of samples tested positive but these were in second round only of nested assay and in some cases only in one of duplicate reactions indicating low levels of virus. From farm 3 (all age groups) for which no mortality events had been reported between 30% to 100% of samples were positive but all in the second round only. On sequencing representatives from all positive farms showed identical sequence (data not shown).

### 3.8 Virus isolation

Fish material from Farms 6 and 7 was inoculated onto BF-2, E-11 and GF cell lines. Cytopathic effects (CPE) were observed in BF-2 and E-11 cells but not GFs on first inoculation (Figure 7). On passage (P1) in the same cell types CPE was only observed for BF-2 cells and intensity of CPE was diminished. Isolated virus from clarified harvested cell culture supernatant from the P1 BF-2 cells was confirmed positive for ISKNV by PCR with sequence identical to that obtained by PCR direct from tissue homogenate (data not shown).

**Figure 7.**
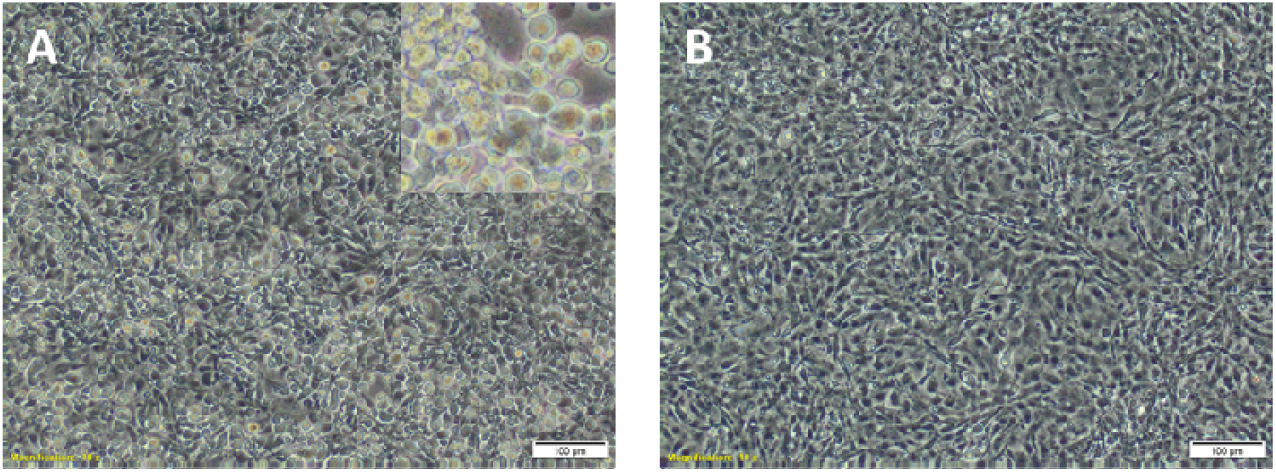
Virus isolation and cytopathic effects in BF-2 cells. (A) BF-2 cells at 5 days post inoculation (−3 dilution) showing enlarged, rounded, phase bright cells. Inset, advanced extensive CPE from −2 dilution of BF-2 at same time point. (B) Negative control BF-2 cells at 5 days.

## 4 Discussion

The results suggest ISKNV had a significant role in the high mortalities experienced by the two farms that were investigated. Fish sampled from the second farm at the height of the disease outbreak showed severe clinical and pathological signs typically associated with infection by the virus (including visualisation of distinctive megalocytes with characteristic virions). Both, these fish and those sampled earlier from Farm 2, had extremely high copy numbers of RSIV/ISKNV *Megalocytivirus-like* virions and ISKNV was confirmed by gene sequencing. Further investigations later in 2019 of five further farms from different areas of lake Volta showed by then the disease was widely established (endemic) in farms across the lake, with both symptomatic and asymptomatic fish positive by PCR for the virus. Farmers emphasised how the disease had a devastating effect on the industry during discussions on this latter visit. They reported how the disease continued to have an impact on the broodstock and growout fish at a lower rate, however the mass mortalities were now predominantly observed in juvenile fish (1-5g in weight). Survival rates to the growout stage, at that time, were estimated to be as low as 5-20%. Juvenile mortalities were reported to be episodic in nature, occurring a few days after the sex reversal process or translocation to lake cages, and lasting up to 3 to 4 weeks. These stress triggered mortality outbreaks may be indicative of either widespread persistence of virus in the environment or a latent ISKNV stage.

To improve survival rates, some farmers had trialled reductions in juvenile stocking densities. Large reductions in stocking density were only associated with small increases in survival, therefore this practice was not considered a viable solution. Instead, farmers resorted to a substantial increase in the level of juvenile production and stocking rates, trading off higher overall mortality with some guarantee of a small but not inconsequential harvest. The economic impact of ISKNV has been significant. The higher production costs and reduced harvests resulted in most farms having to either temporarily or permanently halt production. As larger farms can be the primary employers of some villages, the disease also had a direct impact on the livelihoods of local community members. It was also reported that tilapia market price had more than doubled due to the production shortages and that the feed sales of Raanan Fish Feeds had reduced by 70% (https://thefishsite.com/articles/ghanaian-fish-farmers-seek-help-from-big-business), both potential indicators of the virus having an impact on a much larger scale.

The results would be consistent with a recent introduction of the virus onto one or more farms prior to October 2018, that then extended upstream of the dam to other farms resulting in the unusual, widespread and significant mortalities observed. Firstly, there were no observations of typical ISKNV-associated pathology in any previous disease investigations, on the affected or other farms. Secondly, the limited PCR screening in this study of archived samples of diseased tilapia recovered from before the major mortality event including both farms, were all negative. Moreover, (Verner-Jeffreys et al., 2018) also screened for the presence of RSIV/ISKNV by PCR in 2016, including samples from the affected farms, without detecting the virus, or associated pathology.

Iridioviruses are large icosahedral cytoplasmic double stranded DNA viruses, which can infect a wide range of hosts, including invertebrates and poikilothermic vertebrates. The family *Iridoviridae* includes five genera: *Iridovirus, Chloriridovirus, Ranavirus, Lymphocystivirus* and *Megalocytivirus* (Jancovich et al., 2012). Fish pathogenic iridoviruses are representatives of *Ranavirus, Lymphocystivirus* and *Megalocytivirus* genera (Subramaniam, Shariff, Omar, & Hair-Bejo, 2012; Whittington, Becker, & Dennis, 2010). Infectious spleen and kidney necrosis virus (ISKNV), is a member of the genus *Megalocytivirus* (Jun Kurita & Nakajima, 2012), and causes disease in a range of freshwater and marine fish species (Subramaniam, Shariff, Omar, & Hair-Bejo, 2012; Whittington, Becker, & Dennis, 2010). ISKNV is closely related to red sea bream iridovirus and both viruses are listed by the OIE as notifiable pathogens (World Organisation for Animal Health OIE, 2018). Although tilapia is not listed as a susceptible species by the OIE at present (World Organisation for Animal Health OIE, 2018), recent reports from the USA (Subramaniam et al., 2016) and Thailand (Suebsing et al., 2016) suggest that it is a susceptible species likely to suffer significant mortality. The change in the known host range of virus needs to be communicated to the international community to prevent future transboundary spread through movement of infected tilapia.

It was interesting to note that many of the ISKNV positive fish were actively co-infected with *Streptoccocus agalactiae* and other bacterial pathogens, presenting severe bacteraemia / meningitis, as well as ISKNV-associated pathology. The high mortalities on the farms in the larger on-growing fish may well have been exacerbated by these coinfections.

Although the mortalities were initially confined to on-growing fish in cage culture systems, the later observations of very high ISKNV associated mortalities in fry, associated clinical signs and high viral copy numbers shows the virus likely affects all life stages. As fry are often reared in onshore facilities below the dam and then translocated to on-growing cages on the main lake, this may have been one of the routes that disease was rapidly spread after it first emerged. Anecdotally, at the time of writing, farmers on the sites visited report that mortalities in on-growing facilities have declined, while fry production continues to be badly affected. It is possible that surviving fry have been exposed to the virus and then protected against subsequent exposure. This suggests that immunisation of fry, or use of previously exposed individuals, could represent a practical disease management strategy. Vaccination as a control strategy may be used to control red sea bream iridiodovirus (Nakajima et al., 1999; Shimmoto, Kawai, Ikawa, & Oshima, 2010) and there are also encouraging reports of its potential effectiveness for protection against ISKNV in mandarin fish (Dong, Weng, He, & Dong, 2013). As some of these reports showed efficacy using formalin killed virus infected cells (Dong et al., 2013), rapid development and testing of vaccines based on the direct use of the strain(s) of ISKNV circulating in Lake Volta farms (e.g. autogenous vaccines) should be possible.

Outbreaks of disease that cause significant morbidity and/or mortalities in an aquaculture operation is always a major concern. This is exacerbated when this appears to represent the incursion of a new agent into a system, or region (country or zone in a country) which has not previously been affected. A stark example of this is the epidemic of infectious salmon anaemia virus (ISAV) which reduced production by three quarters and resulted in severe economic and social crisis in the developing Chilean Atlantic salmon industry between 2007 and 2010 (Godoy et al., 2008; Mardones, Martinez-Lopez, Valdes-Donoso, Carpenter, & Perez, 2014; Vike, Stian Nylund, & Nylund, 2009).

The Ghanaian authorities have for some time been concerned that the, to date, successful, expansion of its industry on Lake Volta may be affected by such disease incursions. Partly for this reason, and to safeguard the genetic integrity of Lake Volta Nile tilapia strains, they have tried to limit the culture to locally reared Nile tilapia stocks. However genetic testing by the Ghanaian Fisheries Commission (Ziddah *et al*., Unpublished Observations) showed that fish on some of the farms on Lake Volta were likely of imported GIFT strain origin (Ponzoni et al., 2011) or hybrids of GIFT and indigenous strains.

If farmers have been illegally sourcing broodstock from Asia and other areas, that would be an ideal method of translocating pathogens from one region to another. It should be noted though that ISKNV has also been detected in internationally traded freshwater ornamental species, theoretically posing another possible introduction route (Jung-Schroers et al., 2016)

It is very possible that this is not the first time disease introduction has taken place in Lake Volta Ghana. The study by (Verner-Jeffreys et al., 2018) showed that outbreaks of *S. agalactiae* investigated in 2016 were all caused by genetically indistinguishable isolates of ST 261, with closest genetic identity to Asian isolates. Discussions with affected farmers at the time suggested, that *S. agalactiae* associated mortalities were a relatively recent phenomenon, although the disease is now clearly endemic to all the areas in the Volta area. Other studies have shown that *S. agalactiae* ST261 has likely been translocated around the world in association with farmed tilapia (Kawasaki et al., 2018). As a single large epidemiological unit, it will be difficult to control transmission of virus between farms on lake Volta. It is important to try to prevent spread from Lake Volta to surrounding watersheds and the wider African continent by control over movement of live fish and equipment. The development of biosecure offline hatcheries with borehole water or UV treatment of water, to facilitate production of juveniles which survive to a size they can be vaccinated will likely be key to vaccine control. Additionally, recent technological advances in rapid selective breeding (Houston et al., 2020; Robledo, Palaiokostas, Bargelloni, Martínez, & Houston, 2018) should be employed to develop ISKNV disease resistant populations, or strains of tilapia to enable the industry to recover.

Although most attention to date has focussed on the emergence and spread of TiLV within the tilapia industry world-wide, these results also demonstrate that there is a range of other potential threats to the sustainability of tilapia aquaculture.

## Conclusion

This is the first report of Infectious Spleen and Kidney Necrosis Virus (ISKNV) in farmed tilapia in Africa. ISKNV was found in co-infection with *Streptococcus agalactiae* and other bacterial pathogens in Lake Volta, Ghana. The correlations seen between the mortality events, histopathology and viral loads in the tissues suggest that ISKNV was a major cause of mortalities during the outbreaks. In general, the results support continued efforts to improve the biosecurity of the industry in Ghana. There is a clear need to strengthen domestic capability to rapidly diagnose and control emerging disease threats caused by ISKNV and other pathogens. Further work is also needed to map the distribution of the virus and its impact, including potential effects on wild fish species, and to implement practical control strategies.

## Acknowledgements

The farmers involved in the investigation are thanked for their support and provision of information. Support from Defra (contracts FB002 and FX001 for the OIE Collaborating Centre for Emerging Aquatic Animal Diseases) is gratefully acknowledged.

## Data sharing

The data that support the findings of this study are openly available in BioRxiv doi: https://doi.org/10.1101/680538

