## Supplementary File 1 for "First detection of Infectious Spleen and kidney Necrosis Virus (ISKNV) associated with massive mortalities in farmed tilapia in Africa"

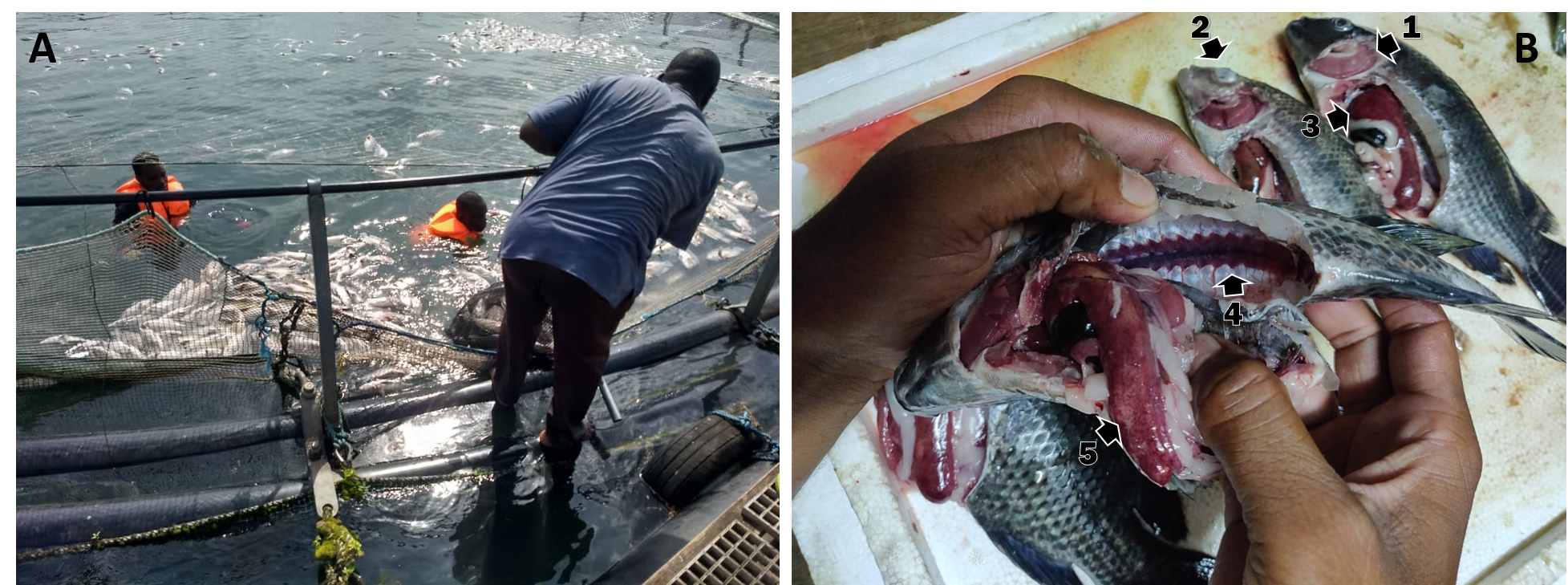

**Supplementary File 1 Mortalities collection and inspection performed by Lake Volta farmers during the outbreaks in October 2018. A** Daily collection by divers. **B** Farmers inspection at farm detecting 1 pale gills; 2 opaque cloudy eye; 3 enlarged haemorrhagic livers; 4 enlarged homorganic kidney; 5 enlarged haemorrhagic liver with white patches.
