## Supplementary File 4 for "First detection of Infectious Spleen and kidney Necrosis Virus (ISKNV) associated with massive mortalities in farmed tilapia in Africa"

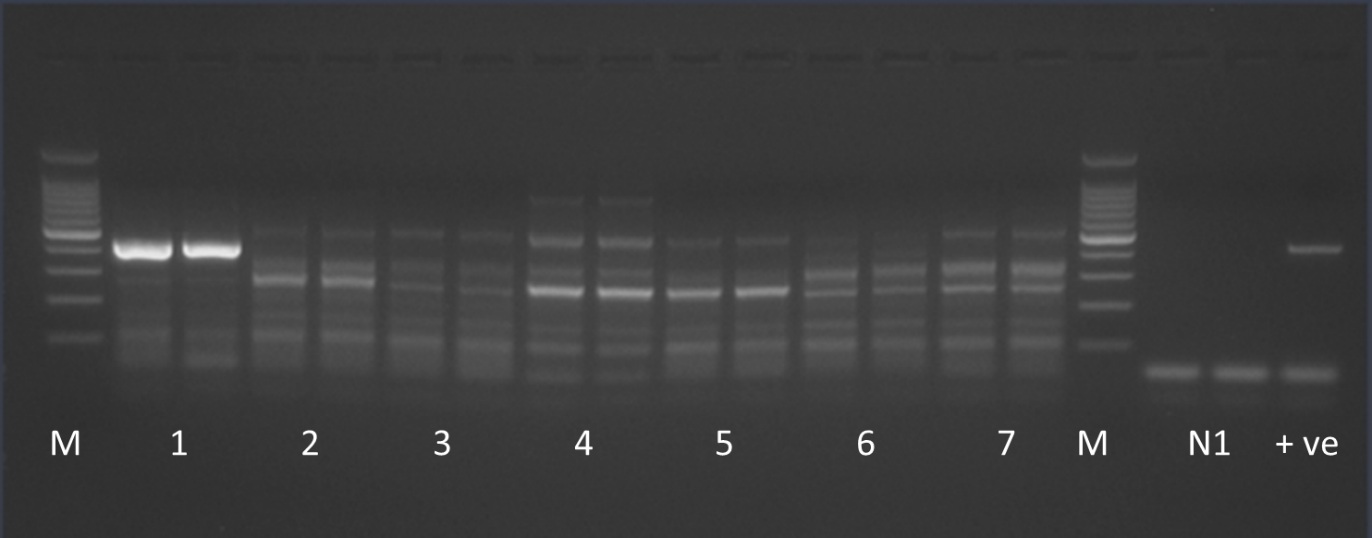
**Supplementary File 4 PCR products generated for the nucleic acid extracted from LV1-7 using the generic *Megalocytivirus* primers.** N1 is the negative control and +ve is the RSIV positive control for the assay. M is the 100bp ladder (Promega).
