## Supplementary File 5 for "First detection of Infectious Spleen and kidney Necrosis Virus (ISKNV) associated with massive mortalities in farmed tilapia in Africa"

| **Supplementary File 5.** Retrospective molecular analysis by qPCR of archived samples. | | | | | |
| --- | --- | --- | --- | --- | --- |
| **Farm** | **Date sampled** | **Organs sampled** | **qPCR_TiLV_** | **qPCR_ISKNV_** | **qPCR_NODA_** |
| Farm 2 | 15/03/2018 | heart, brain and liver | -ve | -ve | -ve |
| Farm 2 | 15/03/2018 | kidney, brain and liver | -ve | -ve | -ve |
| Farm 2 | 15/03/2018 | brain, kidney and liver | -ve | -ve | -ve |
| Farm 1 | 17/03/2018 | spleen, brain, liver and kidney | -ve | -ve | -ve |
| Farm 1 | 17/03/2018 | spleen, brain, liver and kidney | -ve | -ve | -ve |
| Farm 1 | 17/03/2018 | spleen, brain, liver and kidney | -ve | -ve | -ve |
| Farm 3 | 13/03/2018 | kidney, brain and liver | -ve | -ve | -ve |
| Farm 3 | 13/03/2018 | kidney, brain and liver | -ve | -ve | -ve |
| Farm 3 | 13/03/2018 | kidney, brain and liver | -ve | -ve | -ve |
| Farm 2 | 2017 | juveniles <5g | -ve | -ve | -ve |
| Farm 1 | 2017 | juveniles <5g | -ve | -ve | -ve |
| Farm 4 | 2017 | juveniles <5g | -ve | -ve | -ve |
| Farm 4 | 2017 | juveniles <5g | -ve | -ve | -ve |
| Farm 4 | 2017 | juveniles <5g | -ve | -ve | -ve |
| Farm 5 | 2017 | juveniles <5g | -ve | -ve | -ve |
| Farm 5 | 2017 | juveniles <5g | -ve | -ve | -ve |
