## Supplementary File 6 for "First detection of Infectious Spleen and kidney Necrosis Virus (ISKNV) associated with massive mortalities in farmed tilapia in Africa"

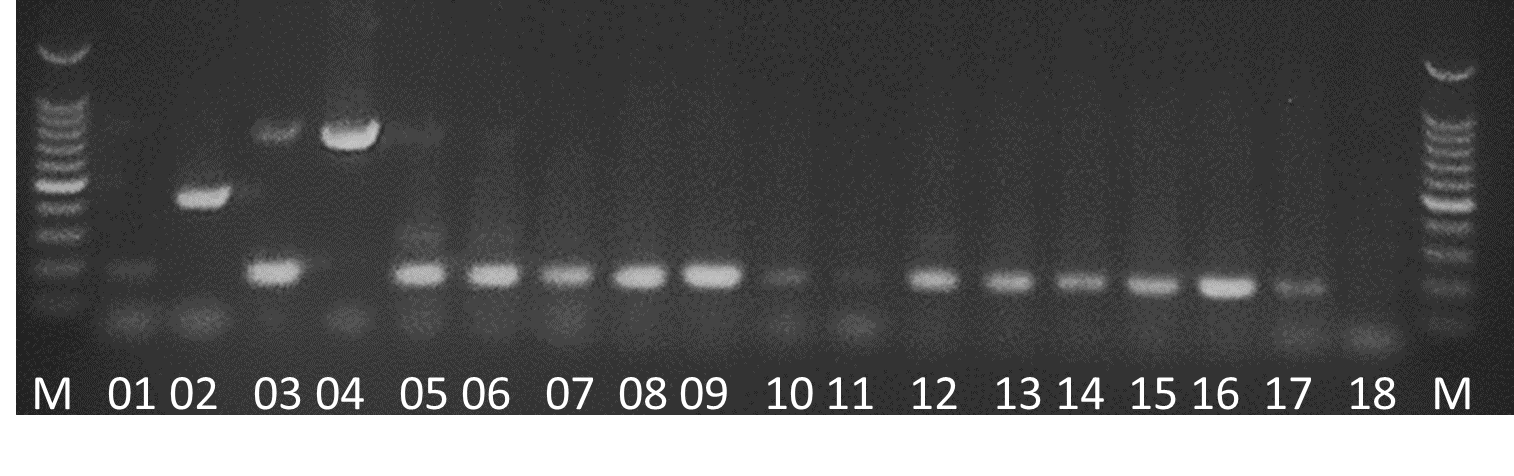

**Supplementary File 6 Multiplex PCR for the detection of 3 different *Streptococcus* spp. in fish tissues.** 01 negative control, 02 *Streptococcus uberis* +ve control (445bp), 03 *Streptococcus agalactiae* +ve control (190bp), 04 *Streptococcus dysgalactiae* (795 bp), 05-18 samples from visit 1.
